# First report of New Clonal groups ST706 and ST1088 from MDR *Klebsiella pneumoniae* Mexican strains

**DOI:** 10.1101/616334

**Authors:** María Guadalupe Córdova-Espinoza, Eder Daniel Hernández Arana, Silvia Giono-Cerezo, Erika Gabriela Sierra Atanacio, Eduardo Carrillo Tapia, Laura Isabel Carrillo Vázquez, Rosa González-Vázquez

**Author notes:** Corresponding authors: PhD. Rosa González Vázquez, Instituto Mexicano del Seguro Social. Hospital General. Centro Médico Nacional. La Raza. Seris y Saachila, SN, Col. La Raza, CP 6450, CDMX, México. Rosa González Vázquez, PhD. María Guadalupe Córdova Espinoza, Departamento de Microbiología. Laboratorio de Bacteriología Médica. Escuela Nacional de Ciencias Biológicas, Instituto Politécnico Nacional, Plan de Ayala y Carpio CP11340, CDMX, México.

## Abstract

Multidrug-resistant *Klebsiella pneumoniae* Mexican strains were characterized for the identification of endemic and pandemic clonal groups. The aims of this study were to know population structure and to identify endemic clonal groups inside *K. pneumoniae* Mexican strains isolated from clinical sources. We studied 93 isolated strains from three third level hospitals and one family clinic from Mexico City. Identification of the strains was done by conventional microbiological methods and an automated system (Vitek2®). The multidrug-resistant phenotype was confirmed following CLSI recommendations, and the strains were classified as MDR, XDR and PDR. Molecular characterization was done by Multilocus Sequence Typing scheme (*rpoB, gapA, mdh, pgi, phoE, infB*, y *tonB*). All strains were isolated from hospitalized patients, the most frequent sources were urine and blood cultures. Population structure of *K. pneumoniae* was clonal, 30 ST were identified, six of them are commonly found. The Clonal complex ST25, ST36, S5392, ST405 and ST551 were isolated from clinical sources, ST1088 was isolated from surfaces of hospital environment.

## Introduction

*Klebsiella pneumoniae* is an opportunist, emerging microorganism, it is an ESKAPE group member (*Enterococcus faecium, Staphylococcus aureus, Klebsiella pneumoniae, Acinetobacter baumannii, Pseudomonas aeruginosa* and *Enterobacter* species). Also, it is responsible of Healthcare Associated Infection (HCAI) (Central Line-associated Bloodstream Infection, Surgical Site Infection, Catheter-associated Urinary Tract Infections, Ventilator-associated Pneumonia) frequently isolated from immunosuppressed patients from intensive care unit (ICU). *K. pneumoniae* is also related to community-associated infections. The most important virulence factors, which allows it to evade the immune response and promote the microorganism’s establishment are: adhesines (fimbriae), capsular polysaccharides (serotypes K1 and K2), siderophores and LPS. (1, 2). *Klebsiella pneumoniae* is frequently identified as a responsible of nosocomial infections. Worldwide, bacterial resistance is an increasing serious problem, as well as extended spectrum beta-lactamases dissemination and different subgroups of CTX-M, which generates cephalosporins resistance, and transferable resistance mechanisms to quinolones and aminoglycosides. The importance of carbapenems transference in plasmids is that they can be transmitted to other bacteria (enterobacteria and other non-fermentative bacilli), as well as the association of resistance to other type of antibiotics. Recent isolation of *K. pneumoniae* KPC (resistance to all beta-lactams, cephalosporins, penicillin and monobactams) at ICU from all over the world is associated to the resistance to other antimicrobial agents, which is contained in the same plasmid (3).

MDR *K. pneumoniae* laboratory identification is done by conventional microbiological methods. Moreover, it’s essential to investigate global dispersion, hospital outbreaks and linage relationships between resistant strains from hospital and community environment, thus, to compare endemic and pandemic clones through molecular typing methods, like Multilocus sequence typing (MLST).

*K. pneumoniae* genetic diversity has been studied previously (4) and it was elucidated that the most dominant ST was from China, other reports show nosocomial clones like ST258, which acquires resistance plasmids with great easiness (5). ST405 is a recently reported clone, which, due to its fast dissemination, is classified as a high-risk clone. The main purpose of this study was to recognize MDR *K. pneumoniae* clones and their prevalence in some Mexican third level healthcare hospitals using MLST technique.

Our results showed that there are clones that were previously reported in distant countries to Mexico. Besides, this paper reports two clones that have not been reported yet. These clones are ST706 and ST1088.

## Materials and Methods

### *K. pneumoniae* strains identification and susceptibility test

93 clinical strains from 2011 to 2013, and 2015 were isolated from different third level healthcare hospital units (hospital I, II, III and IV) from Mexico City. Clinical strains were classified in groups: A, which included outpatient.; I, which was formed by hospitalized patients; and S: inert hospital surfaces isolations. Strains were conserved at 70°C.

*K. pneumoniae* ATCC 700603 and *E. coli* ATCC 25922 were used as controls for the ESBL test (CLSI, 2018) and molecular methods. All isolated clones were identified, and their antimicrobial susceptibilities were analyzed by an automatized system (Vitek2^®^ BioMerieux^®^, France).

Tested antibiotics were: Ampicillin (AMP), cefazolin (CFZ), trimethoprim/sulfamethoxazole (SXT), ampicillin/sulbactam (SAM), cefepime (FEP), ceftriaxone (CRO), aztreonam (ATM), tobramycin (TOB), gentamicin (GEN), nitrofurantoin (NIT), ciprofloxacin (CIP), piperacillin/tazobactam (TZP), amikacin (AMK), ceftazidime (CAZ), levofloxacin (LVX), cefoxitin (FOX), meropenem (MEM), imipenem (IPM) and ertapenem (ETP) (CLSI, 2017). According to susceptibility results, strains were classified as multidrug-resistant (MDR) and extensively drug-resistant (XDR).

For this work, MDR strains were defined as those strains which are resistant to the therapeutic election categories (group A, CLSI 2018). XDR strains were classified as those strains which are resistant to at least one agent in all or two therapeutic election categories (group A and B,CLSI 2018). PDR strains were defined as those strains resistant to all agents in all antimicrobial categories (group A, B, C and U, CLSI 2018) however colistin was not taken for resistance classification because its associated with nephrotoxic damage and it is not commonly used in clinical hospitals. Clinical isolates with resistance to one antibiotic in less than three categories were considered as not classifiable.

### Confirmatory Extended-Spectrum β-lactamases by phenotype detection test

ESBL confirmative test was performed using the Double Disc Synergy Test employing ceftazidime (30 μg), ceftazidime-clavulanate (30 μg/10 μg), cefotaxime (30 μg), and cefotaxime-clavulanate (30 μg/10 μg) disks. Quality control was done using *K. pneumoniae* ATCC 700603 and *E. coli* ATCC 25922. ≥ 5mm increase in diameter of ceftazidime-clavulanate or cefotaxime-clavulanate indicates an ESLB producing microorganism (positive result) (CLSI, 2018).

### Molecular characterization

48 strains were selected to perform MLST scheme, strains selection was done considering hospital unit, clinical sample origin, isolation year and results from susceptibility test. DNA was extracted by guanidine thiocyanate method. Multilocus sequence typing scheme (*gapA, infB, mdh, pgi, phoE, ropB* and *tonB*) has been described before by Diancourt (6). DNA PCR products were purified by PureLink Quick gel extraction (ThermoFisher Scientific, USA), PCR purification combo Kit (Invitrogen, USA) and EZ-10 Spin Column PCR purification Kit (BioBasic, Canada). Sequencing PCR protocol was done using ABI PRISM Big dye Termination v3.1 Cycle Sequencing Kits. ABIPRISM 3730XL equipment was employed to read sequencing reactions.

### Sequence data analysis

Blast alignment for each housekeeping gene was done to verify the sequence identity (http://www.ncbi.nlm.nih.gov/). FinchTV v 1.4.0 (FinchTV 1.4.0 Geospiza), ClustalX v 2.1 (7) and Bioedit v 7.2.5 (8) programs were used to edit genes sequences. Finally, sequences were submitted to *K. pneumoniae* PUBMLST web site (“http://pubmlst.org/”) and gene alleles and sequence type (ST) were assigned.

eBURST v3 program was used to obtain clonal complex (CC) (9). Sequence type analysis was done with eBURST v3, START2 v 0.9.0, and DNAsp v5.10 program (10, 11). ExPASY (www.expasy.org) web site was also used for amino acid translation. Phylogenetic networks were constructed using START2 v 1.0.5 (http://pubmlst.org/software/analysis/start2/) program, *Neighbor-joining* method and *balanced minimum evolution* (BME) criteria.

## Results

### *K. pneumoniae* strains identification and susceptibility test

Ninety-three *K. pneumoniae* strains were isolated from different clinical samples, the 49.5% (46/93) were isolated from urine, 21.5% (20/93) from blood, 9.7% (9/93) from hospital surfaces, 8.6% (8/93) from catheter tip, 8.6% (8/93) from organic liquids and 5.4% (5/93) from respiratory secretions.

Antimicrobial resistances obtained were as follows: AMP, 96% (89/93), CFZ, 61% (57/93), SXT, 60% (56/93), SAM, 57% (53/93), FEP, 56% (52/93), CRO, 56% (52/93), TOB, 39% (36/93), GEN, 35% (33/93), NIT, 27% (25/93), CIP, 22% (20/93), TZP, 16% (15/93), AMK, 9% (8/93), MEM, ETP 2% (2/93) and IPMM, 1% (1/93). Sixty percent of the population (56/93) were classified as ESBL producing strains by Vitek2^®^. The results were confirmed by confirmatory test and they were matched with the Vitek2^®^ results. Classification resistance tests allowed us to distinguish 46% of the population (43/93) as MDR strains, 17% (16/93) as XDR strains, and 1% (1/93) as PDR strains. 36% of the population (33/93) were not classifiable.

### MLST results

Phylogenetic analysis was performed with 48 strains and 2317 ST previously reported in the MLST data base (http://bigsdb.pasteur.fr/). In this study, 30 ST were identified:29 ST corresponding to clinical strains and 1 ST from *K. pneumoniae* ATCC 700603 (table S1). Burst algorithm identified groups of related genotypes (clonal complex) and allowed to recognize the founding genotype of each group, though, analysis population snapshots were obtained and showed a clonal structure [I_A_ = 0.572 (n=48) (START2 V.1.0.5)].

eBURST v3 program considers the 30 ST (including ATCC 700603 strain with ST489, circled in green; and the clinical isolated strains, circled in red) that were identified in this study with repetitions (1000 “bootstrap”). CC denomination was assigned according to each group’s founding ST, and it is considered that a CC is composed by isolations that have 6/7 identical loci. There was a total of 183 generated CC, the principal clonal complex (CC11) is observed in the center of the “snapshots”, this complex showed 47 SLV, 43 DLV, 123 TLV and 688 SLV, in where 18 ST’s were found. Following CC to which the isolated strains belong are: CC147 (ST885, ST392); CC628 (ST628); CC1088 (ST1088); CC551 (ST551); CC2054 (ST10726); CC491 (ST491); CC405 (ST405) and CC307 (ST307). Individual ST’s (“singleton”) that were not part of a CC were ST2080, ST1846 and ST489. 23 ST’s (79%) were from 1 strain, 6 ST (21%) were from 2 or 6 isolates. Predefined ST were ST551 and ST405 (12.5%, 6/48), then ST25 and ST1088 (8.3%), right away ST392 (6.3%, 3/48%) and finally ST36 (4.1% 2/48). The six ST were considered the clones prevalent in this study. MDR strains were ST25 (2/4), ST405 (2/6) (Fig. S1).

Isolations obtained were classified as follows: 46% (43/93) were MDR, 24 from Hospital I, 10 from Hospital II, 1 from Hospital III and 8 from hospital IV; 17% (16/93) were XDR, 10 from Hospital I, 1 from Hospital II, 2 and 3 from Hospital III and IV. PDR strain (1%, 1/93) was isolated form Hospital IV. ST392 (4/4), ST 1088 (3/3) and ST551 (3/6); XDR strains were ST25 (2/4), ST551 (3/6) and ST405 (2/6); ST706 were PDR.

### Sequence Analysis

Characteristics and polymorphism of each gen are shown on Table S4*. Each gen number of alleles varied from 4 (*gapA* y *rpoB*) to 17 (*tonB*), and the number of polymorphic sites match with each gene mutations. Values of π, from 0.00463 (*pgi*) to 0.02794 (*gapA*) and θ from 0.00364 (*gapA*) to 0.01331 (*rpoB*) were < 1. G+C content was found over the 50% in the seven genes, with ranges from 55.77% (mdh) to 64.66% (tonB).

Polymorphic changes match with the mutations, most of the “housekeeping” genes did not show mutations on the different triplet positions, with synonymic and non-synonymic amino acid changes. *dn* / *ds* relation for most genes were significatively <1, indicating that there was not a strong selective pressure over the genes, thus why these genes do not affect bacterial viability, excepting *infB* gene.

### Phylogenetical Relation

Phylogenetical analysis was done with 48 isolated strains, following “*Neighbor-joining*” method, based on BME (“*balanced minimum evolution*”) criteria, which determines the closest sequences by binding them with an internal node at an 0.1 distance, repeating itself on the remaining sequences until all of them are linked by those internal nodes, which minimize each internal branch length, thus obtaining a phylogenetical tree in which its branches length indicates de evolutive changes (Figure S3).

The phylogenetical tree shows an extern group, conformed by 10% (5/48) of isolated strains, each with a different ST (ST846, ST491, ST111, ST804 y ST16), whereby, phylogenetical distance varied. Internal group was divided into two different clads (aggrupation with a common predecessor), A clad includes 50% (24/48) of the isolated strains, while B clad includes 40% (19/48). Strains with the same ST (ST392, ST551, ST36, ST1088, ST405, ST25) are found in the same clad and at the same distance. There was no observed relation between antibiotic resistance with phylogenetical groups formation.

## Discussion

According with the CDC latest data, ESKAPE group bacteria are held responsible of two thirds of all the Healthcare Associated Infections and play a significant role in worldwide mortality. *Klebsiella pneumoniae* belongs to this groups. In Mexico, *Klebsiella pneumoniae* is classified as one of the three principal HCAI etiologic agents, as well as the second most reported microorganism in outbreaks. In the last 5 years, there has been an increase on the number of cases. Though it is saprophytic bacterium found in gastrointestinal tract, skin, nasopharynx, it can also cause community and hospital infections with a lethality rate of 35% (12).

In this study, ninety-three *K. pneumoniae* isolations, coming from 4 different hospitals located on Mexico City were analyzed. The most frequent isolation source was urine culture followed by blood culture. The RHOVE 2016 report emphasizes that *K. pneumoniae* got second place in isolation frequency in bloodstream infections, and fourth place in urinary system infections. There have been reports in other countries about high-frequency isolations in blood and urinary system infections in third level hospitals contrary to the reported by Shanmuga & Usha (2018) were bronchial secretion samples were found in a low frequency.

On 2017, the WHO published for the first time the priority pathogen list, such pathogens represent a public-health threat, considering the antibiotic resistance. Carbapenem resistant, EEBL producer *K. pneumoniae* is included in “critical priority” (14).

Gashaw performed a study in a third level hospital, obtaining 30% MDR, 43% XDR and 7% PDR. β-lactam resistance is an increasing problem for the infection treatment, thereby, use of carbapenem antibiotics for treatment of infections caused by EEBL producing microorganisms was implemented. Nevertheless, carbapenem resistant strains have been reported, elevating patients’ mortality index (15).

Antibiotic resistance was determined in clinical isolated strains using Vitek2^®^, so classification according to criteria described in this work, could be applied, obtaining a higher MDR strains percentage (16) reported that 80% of a Mexico City hospital isolated strains were MDR, nevertheless, no PDR strain was obtained, whereas in this study, one PDR strain was found. This allows us to visualize the increasing resistance of *K. pneumoniae* due to indiscriminate use of antibiotics for treatment of infection caused by this bacterium. Resistance is associated with the expression of genes contained in plasmids, transposons and integrones which are key elements in horizontal genetic material transference. *Klebsiella* carbapenem resistance has been principally associated with plasmids (15, 17).

On this work, MLST technique was applied for the establishment of a genetic relation on clinical isolated strains associated to outbreaks of *K. pneumoniae*. 48 isolations were analyzed, beside a control strain of K. pneumoniae ATCC 700603 used for validation, obtaining ST489 for such strain (18). On the clinical isolated strain analysis, 29 type sequences were obtained and none of the new alleles contained on the PubMLST database. 23 ST matched to one strain, and the six remaining strains were considered as clones (ST551, ST405, ST25, ST1088, ST392, ST36). Diverse clones had been previously reported on distant countries from Mexico, which allows us to visualize the rapid dissemination of some clones.

Index association was done for the present clones in this study, allowing to confirm with eBRUST algorithm that the studied population was clonal type, in other words, the isolated strains are genetically related.

“Neighbor-joining” method was used for the construction of a phylogenetical tree and for a better visualization of the phylogeny analysis. The analysis showed two principal clads. The A clad contains 50% of the isolations coming from the 4 mentioned hospitals, of these, 13 were classified as MDR, 5 as XDR and 1 as PDR. Predominant ST’s for this clad were: **ST551**, isolated on 2012 from hospital I and from hospital II on 2013 from patients and surface samples. This strain was held responsible for an outbreak on Azcapotzalco delegation on Mexico City. This clone was also isolated on years 2011 – 2013 in Japan, according to (19), where its major isolation frequencies were in urine and respiratory secretion samples.

**ST392** was found on hospital IV on 2015, this strain was classified as endemic for the hospital. **ST392**, was detected during an 11-month survey; also, **ST307** (20), reported as well in Italy (21), where they mention that this clone is highly risky due to its rapid dissemination, plus the characterized isolations of this strain were KPC producers. This specific clone has been reported in Korea, Pakistan, Italy, Morocco, Mexico, Serbia and Japan. Because it’s been reported in different countries worldwide, this clone is a strong candidate to become a high-risk clone in a near future.

**ST36** was isolated from hospital I on 2015 and from hospital II on 2013; all isolations were susceptible to previously mentioned antibiotics, meaning an advantage por the patient’s treatment, because the bacteria can be dealt with following the medical instructions, surface sanitization and adequate treatment. Nevertheless, on countries like China, this same clone has been reported as a hyper virulent strain with carbapenem resistance (22). **ST706** was the only PDR strain, and was isolated from hospital IV on 2015. There are no current published papers about this strain.

B clad contains 39.6% of the isolations, 10 were classified as MDR and 4 as XDR, according to the classification. Predominant ST’s in this clad were: **ST405** in hospital I, correspondent to years 2011, 2012 and 2015; in hospital II on 2013 and in hospital IV in 2015. Its most frequent isolation sources were urine and blood culture.

**ST405** clone was responsible for outbreaks in Spain in years 2010-2012, nevertheless, it was disseminated time after that to countries like Italy, where it caused an outbreak on years 2013 – 2014, later to Austria and Germany. Studies performed on 2019 mention that, unlike the isolations obtained in Spain, these isolations did not present OXA-84 (23). Nevertheless, they presented other virulence factors. This was attributed to the fact that most of the samples obtained were taken from water bodies near the hospitals, better said, community waters. These strains have proved susceptible to antibiotics, while hospital strains are resistant, but wild strains express more virulence factors.

**ST25** clone was found in hospitals I and IV on 2015, and in hospital II on 2013. This clone was isolated in Japan on 2008, and in China on the June 2014-May 2015 period. ST1088 was found only on inert surfaces in hospital II on 2013 and was classified as endemic for such hospital. This suggests that no adequate survey or preventive measures for this bacterium have been taken, that is why it is important to rotate the disinfecting solutions every once I a while.

These ST’s present major importance since they’re present in at least three hospitals located in different areas inside Mexico City.

ST258 is one of the most widely disseminated and most prevalent clones, there are several reports about this clone presenting resistance to multiple pharmaceutics, including carbapenems. Nevertheless, this clone was not found in this study. There have been reports about carbapenem resistance associated clones in Brazil and China, such as ST11, ST25 and ST392. These clones were obtained in this study but showed sensibility to these antibiotics. However, precaution must be taken because these strains can easily obtain genetic material for carbapenem resistance.

ST’s reported in this study had been previously reported in other countries’ outbreaks, except **ST706** and **ST1088**, however, in Mexico, not all the reported clones were well descripted. That’s why it is important to emphasize the importance of the survey, since great part of them were classified as MDR. This allows us to evaluate the status of *K. pneumoniae* caused infections in Mexico by comparing it with previously published studies, observing the increase in resistance and dissemination of diverse clones that were reported in other countries.

## Acknowledgments

This work’s authors thank Dr. Laura López Pelcastre and C. Carmen Melchor Díaz, as well as the Mexican Institute for the Social Security for their input into this project.

---

**FIG S1.**
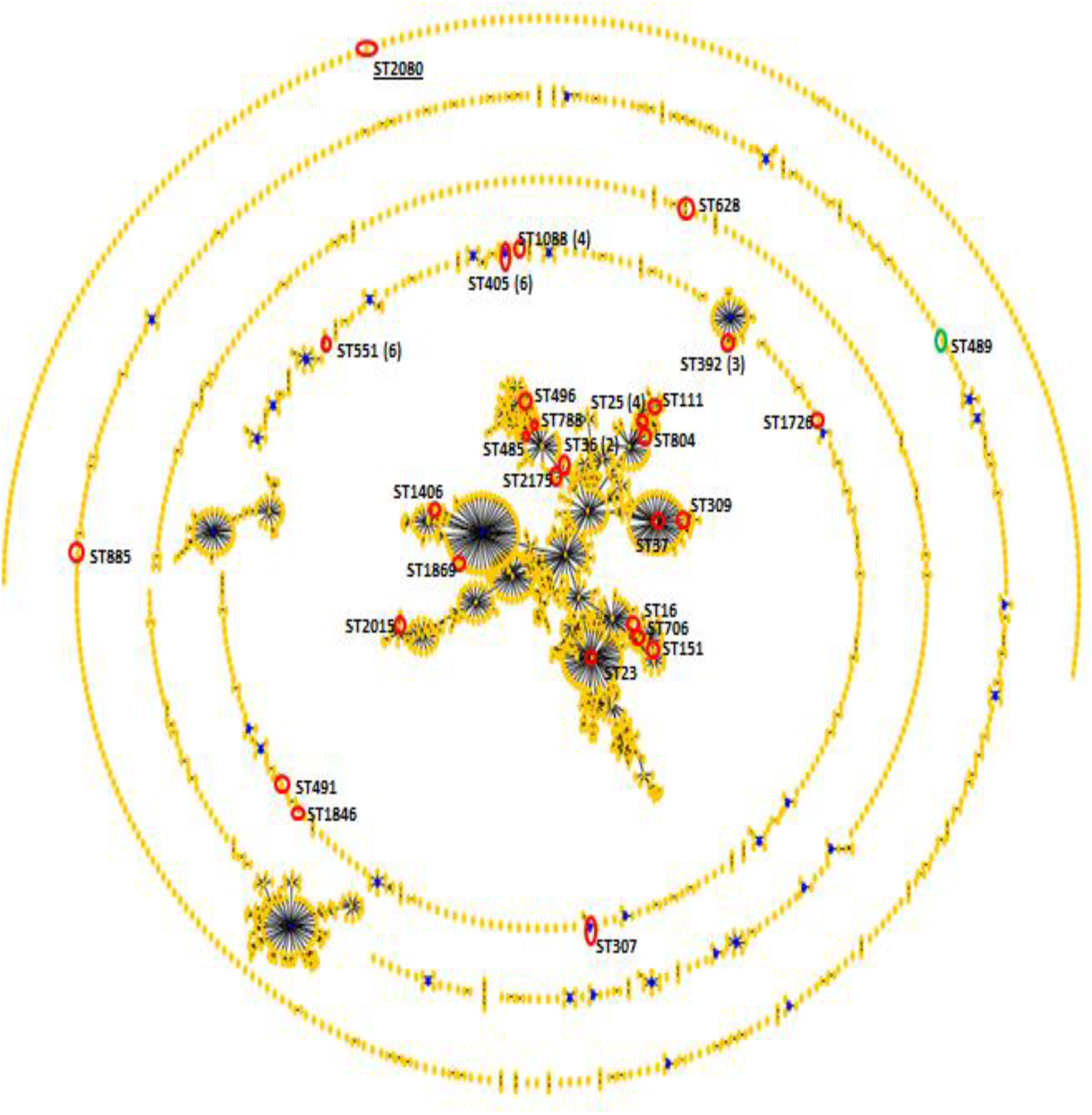
Snapshot from *K. pneumoniae*, comparative analysis eBURST. 48 ST from *K.pneumoniae* were identified (red circle); *K. pneumoniae* ATCC 700603 (green circle); founders were in the center of each clonal complex. Individual ST were highlighted with a a line.

**Table S2.**
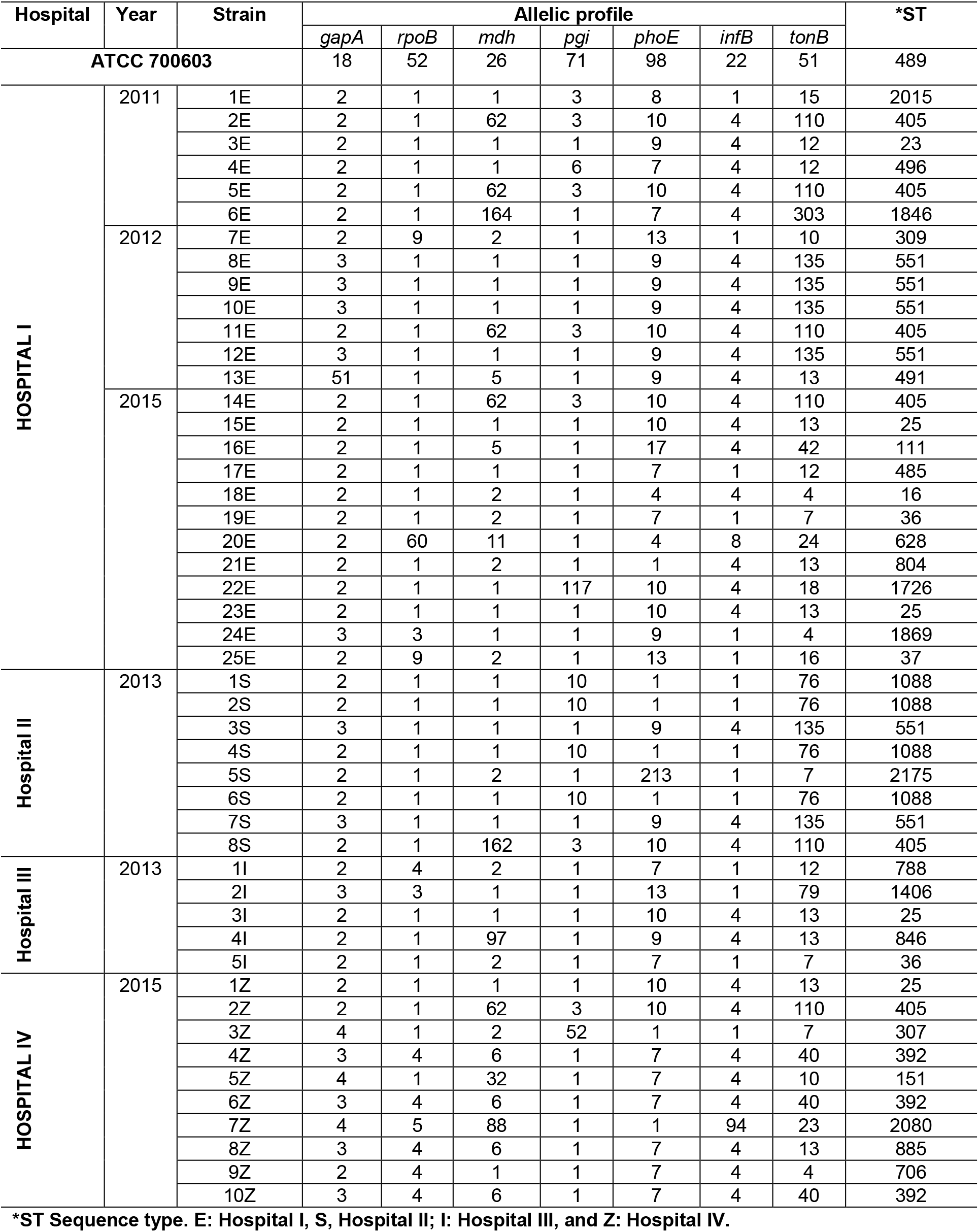
Allelic form gene and ST assignment from clinical and reference strains

**FIG S3.**
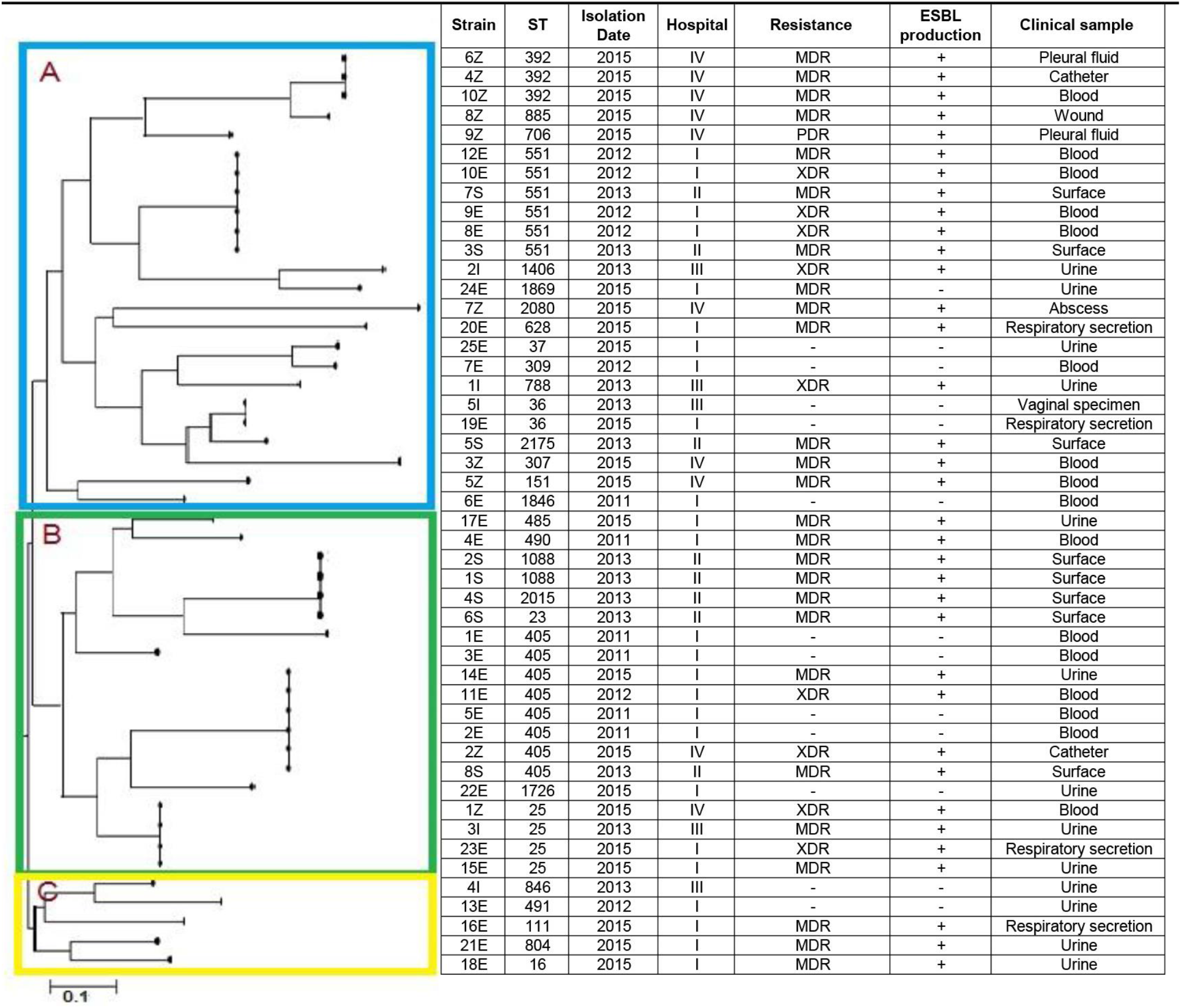
Phylogenetic tree from clinical samples of K. pneumoniae strains obtained by Neighbor-joining method, through the START v.0.9.0 program. The length of the branches of the phylogenetic tree indicate the evolutionary change of each isolate. A and B form the intern group, each one them represent a clade and subdivide on subclades. C is a extern group.

**Table S4.** Characteristics and polymorphism of the *housekeeping* genes from K. pneumoniae isolated.

| Gene | PCR product size | Haplotype | Polymorphic sites | Total mutations | $\pi^*$ | $\theta^*$ | G+C | dN* | dS* | dN/dS |
| --- | --- | --- | --- | --- | --- | --- | --- | --- | --- | --- |
| *gapA* | 450 | 4 | 3 | 3 | 0.02794 | 0.00364 | 0.5615 | 0.0000 | 0.0132 | 0.0000 |
| *infB* | 318 | 6 | 7 | 7 | 0.00797 | 0.00964 | 0.6140 | 0.0097 | 0.0037 | 2.6548 |
| *mdh* | 477 | 10 | 21 | 21 | 0.00526 | 0.00684 | 0.5577 | 0.0086 | 0.0145 | 0.5944 |
| *pgi* | 432 | 6 | 6 | 6 | 0.00463 | 0.00608 | 0.5733 | 0.0042 | 0.0056 | 0.7531 |
| *phoE* | 420 | 9 | 9 | 9 | 0.00728 | 0.00788 | 0.5585 | 0.0000 | 0.0311 | 0.0000 |
| *rpoB* | 501 | 4 | 13 | 13 | 0.01331 | 0.01331 | 0.5415 | 0.0128 | 0.0147 | 0.8713 |
| *tonB* | 414 | 17 | 16 | 16 | 0.01027 | 0.01143 | 0.6466 | 0.0050 | 0.0257 | 0.1961 |

$\pi^*$ : Nucleotide diversity per site.
$\theta^*$ : Average of nucleotide differences per site.
dN\*: Numbers of non-synonymous substitutions per site.
dS\*: Numbers of synonymous substitutions per site.

